# Brassinosteroid signaling component SlBES1 promotes tomato fruit softening through transcriptional repression of *PMEU1*

**DOI:** 10.1101/2021.01.18.427052

**Authors:** Haoran Liu, Lihong Liu, Dongyi Liang, Min Zhang, Chengguo Jia, Mingfang Qi, Yuanyuan Liu, Zhiyong Shao, Fanliang Meng, Songshen Hu, Chuanyou Li, Qiaomei Wang

## Abstract

Firmness is one of the most important factors that affect postharvest properties of tomato fruit. However, the regulatory mechanism underlying firmness formation in tomato fruit is poorly understood. Here, we report a novel role of SlBES1, a transcriptional factor (TF) mediating brassinosteroid (BR) signaling, in tomato fruit softening. We first found that SlBES1 promotes fruit softening during tomato fruit ripening and postharvest storage. RNA-seq analysis suggested that *PMEU1*, which encodes a pectin de-methylesterification protein, might participate in SlBES1-mediated fruit softening. Biochemical and immunofluorescence assays in *SlBES1* transgenic fruits indicated that SlBES1 inhibited *PMEU1*-related pectin de-methylesterification. Further molecular and genetic evidence verified that SlBES1 directly binds to the E-box in the promoter of *PMEU1* to repress its expression, leading to the softening of the tomato fruits. Loss-of-function *SlBES1* mutant generated by CRISPR/cas9 showed firmer fruits and longer shelf life during postharvest storage without the color, size and nutritional quality alteration. Collectively, our results indicated the potential of manipulating *SlBES1* to regulate fruit firmness via transcriptional inhibition of *PMEU1* without negative consequence on visual and nutrition quality.

## Introduction

Brassinosteroids (BRs) are well-characterized phytohormone that are critical for plant vegetative growth, development, and response to environmental stimulus, influencing many important agronomic traits (He et al., 2005; Yu et al., 2011; Chen et al., 2019; Cui et al., 2019; Nolan et al., 2020). Recently, BRs were found to be actively synthesized and accumulated during tomato fruit ripening, indicating an important role of BR signaling in fruit (Li et al., 2016; Hu et al., 2020). Thus, the role of BRs in the regulation of fruit ripening remains to be further investigated considering the potential agricultural and research value. Genetic and molecular studies in *Arabidopsis* have revealed many core components of BRs signaling pathway, which constitutes a complete signal pathway from cell surface BR receptor to downstream transcription factors that regulate the expression of BR-regulated genes (Ye et al. 2011; Guo et al. 2013). BRI1 -EMS-SUPPRESSOR1 (BES1), a key basic helix-loop-helix TF in the BR signaling pathway, balances plant growth and environment stress tolerance (Yu et al., 2011; Nolan et al., 2020) via binding a conserved E-box (CANNTG) and BRRE (CGTGC/TG) elements of its target genes (Cui et al., 2019; Jiang et al., 2019). Although the mechanism of BES1 transcription factor in regulating plant growth and development has been well elucidated in *Arabidopsis*, its function in fruit softening has remained elusive.

Tomato (*Solanum lycopersicum*) is not only one of the most important vegetables worldwide, but also a model system for researchers studying fruit ripening. The softening process during ripening indicated by firmness determines overall palatability, transportability, and shelf life of tomato fruit. Fruit firmness is determined by diverse factors, including cell wall structure (Seymour et al., 2013), cell turgor (Saladié et al., 2007), and cuticle properties (Yeats and Rose, 2013). Among them, cell wall and alterations of its structure are considered as the predominant factors, such as changes in the complex of microfibrils and polysaccharides, which are composed of hemicellulose, cellulose, pectin, and other structural proteins. Pectins, known as pectic polysaccharides, are a group of complex polymers containing homogalacturonan (HG), rhamnogalacturonan-I, and rhamnogalacturonan-II and located in the primary cell wall and middle lamella with a high amount (Brummell, 2006). A wide range of enzymes have been investigated to explore the genetic, molecular and biochemical basis of fruit softening. Methyl ester group is removed from HG in a reaction catalyzed by pectin methylesterase (PME/PE, EC 3.1.1.11), and the de-methylesterified pectin forms “egg box” through a cross-link with Ca^2+^, strengthening the cell wall (Senechal et al., 2014; Silva-Sanzana et al., 2019). In addition, pectin degradation by pectate lyase (PL) and polygalacturonase (PG) were paid attention to targeted control of fruit softening (Uluisik et al., 2016; Wang et al., 2019). Nowadays, tomato cultivars harboring *ripening inhibitor* (*rin*) and *non-ripening* (*nor*) mutation produce firm fruits and confer long shelf life. However, they often have a deficiency in coloration and poor nutritional value because of the disturbed ethylene-regulated fruit ripening (Klee and Giovannoni, 2011; Osorio et al., 2011; Osorio et al., 2020). Therefore, targeted control of fruit softening without negative consequences on fruit quality by dissecting the regulatory mechanism becomes an important goal to extend shelf life and improve transportability.

In this study, the specific role of SlBES1 in softening was investigated in tomato. Firmness phenotypes of *SlBES1*-overexpressing and silencing fruits indicated its positive role in softening. RNA sequencing (RNA-seq) in silencing fruits elucidated the possible target genes, including *pectin methylesterase ubiquitously 1* (*PMEU1*), responsible for softening. More biochemical, molecular and genetic experiments further reveal that SlBES1 inhibited PMEU1-associated pectin metabolic pathway and directly mediated transcriptional regulation of target gene involved in softening. Interestingly, knockout of *SlBES1* in tomato prolonged shelf life without detrimental effects on nutrition quality, implying SlBES1 could be a potential powerful biotechnological tool in tomato breeding for improvement of shelf life.

## Results and discussion

### Functional identification of SlBES1 in tomato

The central role of BES1 in mediating growth and stress response has been widely investigated, but few studies focused on its role in fruit development. To verify the function of BES1 in fruit development, we identified *SlBES1*, the homologous gene of *AtBES1* in tomato (Supplemental Figure S1A and B), and analyzed its expression pattern in BR-deficient and BR-insensitive tomato mutants. The expression level of *SlBES1* was decreased in both BR-deficient mutant *d^x^* and BR-insensitive mutant *cu-3* that harbors a mutation in the BR receptor gene *BRI1* (Supplemental Figure S1C). To explore the function of SlBES1 in tomato, we then generated *SlBES1* overexpressing transgenic lines, *SlBES1-OX-3* and *SlBES1-OX-8*, as well as *SlBES1* RNAi lines, *SlBES1-RNAi-8* and *SlBES1-RNAi-9* (Figure 1A). In *Arabidopsis*, the gain-of-function mutant of *AtBES1* (*bes1-D*) and *AtBZR1* (*bzr1-D*) have constitutive BR responses including tolerance to BR biosynthesis inhibitors, propiconazole (Pcz) or brassinazole (BRZ), in hypocotyl elongation assays in the dark (Yin et al., 2002; Wang et al., 2002; Hartwig et al., 2012). The hypocotyl elongation of *SlBES1-OX-3* and *SlBES1-OX-8* was not affected by 0.5 μM Pcz in the dark, whereas that of *SlBES1-RNAi-8* and *SlBES1-RNAi-9* was more susceptible to inhibitor (Supplemental Figure S2A and B), which was in agreement with the phenotypes of *Arabidopsis BES1* mutants. In addition, BES1/BZR1 repressed the transcription of BR biosynthetic genes, such as *DWARF*, *CPD*, and *DWARF4*, through feedback regulation (He et al., 2005; Yu et al., 2011). In the present study, expression levels of the BR biosynthetic genes (*SlDWARF*, *SlCPD*, *SlCYP724B2*, and *SlCYP90B3*) were significantly down-regulated in *SlBES1-OX-3* and *SlBES1-OX-8*, but up-regulated in *SlBES1-RNAi-8* and *SlBES1-RNAi-9* fruits (Supplemental Figure S2C). These results indicate that SlBES1 confers conserved function as a transcription factor in BR signaling pathway between tomato and *Arabidopsis*.

**Figure 1.**
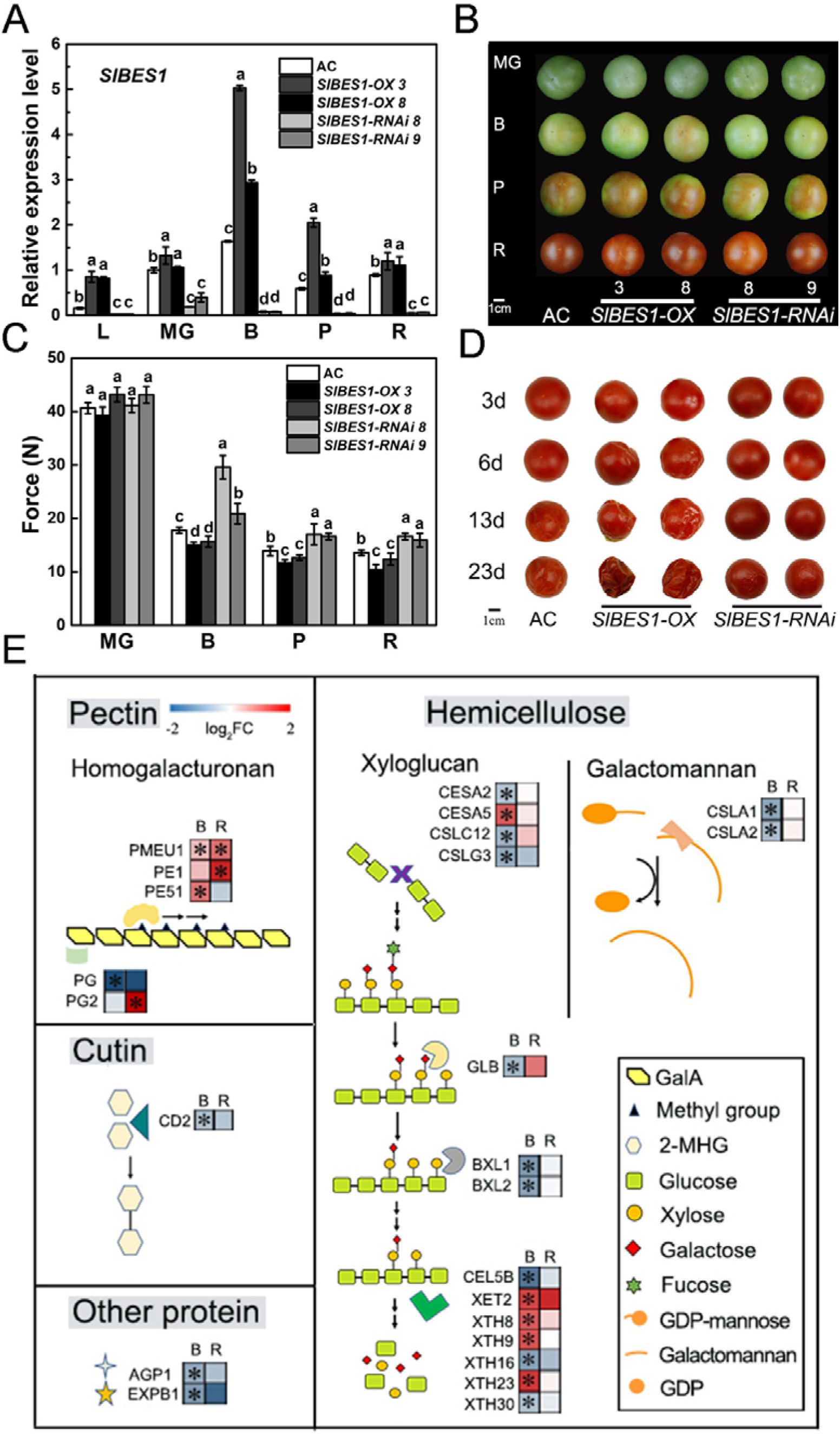
Phenotypes of *SlBES1-OX* and *SlBES1-RNAi* Tomato Fruits. (A) Gene expression levels of *SlBES1* in leaves and fruits of *SlBES1-OX* and *SlBES1-RNAi* plants. L, leaves. MG, mature green stage. B, breaker stage. P, pink stage. R, red ripe stage. Data shown represent the means±SD of three biological replicates. Different letters indicate significant difference compared to AC (wild type) (one-way ANOVA with Tukey’s test, *p*<0.05). (B) Phenotypes of AC (wild type), *SlBES1-OX* and *SlBES1-RNAi* tomato fruits. Pictures were taken at four developmental stages of fruit. MG, mature green; B, breaker; P, pink; R, red ripe. Scale bar = 1 cm. (C) Fruit firmness of AC (wild type), *SlBES1-OX* and *SlBES1-RNAi* fruits at different development stages. MG, mature green. B, breaker. P, pink. R, red ripe. Data shown represent the means±SD of twenty-four biological replicates from at least six independent fruits. Different letters indicate significant difference compared to AC (wild type) (one-way ANOVA with Tukey’s test, *p*<0.05). (D) Tomato shelf life of AC (wild type), *SlBES1-OX* and *SlBES1-RNAi* fruits. Fruits were harvested at red ripe stage and stored at room temperature. The progression of fruit deterioration was recorded by time-lapse photography. Time after harvest is specified by days. The storage condition was 25°C and 35% relative humidity. Scale bar = 1 cm. (E) RNA-seq results and related metabolic pathway of *SlBES1-RNAi* fruits at B and R stages. A heat map showing the expression pattern of firmness related genes in *SlBES1-RNAi* fruits and wild type. The values of fold change (log_2_) with or without significance (*p*<0.05) were represented as coloured block with or without asterisk, respectively. Each group contains three biological replicates. B, breaker stage. R, red ripe stage. Accession number, name, description and the value of fold change of genes in figure were listed in Supplemental Table 1.

### SlBES1 promotes tomato fruit softening without affecting nutritional quality

Transgenic *SlBES1* fruits were used to elucidate the role of BES1 in fruit development. As shown in Figure 1B, no significant difference in fruit appearance at different developmental stages was found between wild type Ailsa Craig (AC) and transgenic tomato lines. However, *SlBES1-OX* lines and *SlBES1-RNAi* lines exhibited reduced and increased fruit firmness compared with AC, respectively (Figure 1C). These varying degrees of firmness correspond to the *SlBES1* expression levels in different transgenic lines (Figure 1A), which demonstrated that SlBES1 negatively regulates tomato fruit firmness. The production of ethylene in *SlBES1* transgenic tomato fruits and AC was similar at each development stage (Supplemental Figure S3), indicating SlBES1-mediated softening is via ethylene-independent manner and different with firmer but ripening-impaired mutants, *rin* and *nor*. Meanwhile, overexpressing or silencing of *SlBES1* did not affect nutritional qualities (Supplemental Table 3) and the expression levels of most nutrition related genes, including carotenoid and ascorbic acid biosynthetic genes, in *SlBES1-RNAi* kept at the same level with wild type (Supplemental Table 4).

Furthermore, treatment of wild type AC fruits with 3 μM 24-epibrassinolide (EBL) significantly promoted fruit softening (Supplemental Figure S4A). Meanwhile, EBL treatment recovered fruit softening in the BR-deficient mutant *d^im^* (Supplemental Figure S4B), indicating the role of BR in cell wall homeostasis. Fruit firmness is usually essential in determining fruit shelf life. We also detected a complete loss of fruit integrity and shorter shelf life in *SlBES1-OX* but longer shelf life in *SlBES1-RNAi* when compared with that of the wild type (Figure 1D). The *SlBES1-OX* lines have similar phenotype as the loss of integrity in fruits with inhibited PME activity (Tieman et al., 1992; Tieman and Handa, 1994).

To reveal firmness-related genes or metabolism processes that participate in SlBES1-induced fruit softening, *SlBES1-RNAi* fruit was harvested at the breaker (B) and red ripe (R) stages for RNA-seq. Gene ontology (GO) analysis on RNA-seq data in fruit development period category (*p* < 0.05) showed that the expression of 24 genes related to fruit firmness, including the pectin metabolism gene *PME* and hemicellulose metabolism gene *XTH*, was differentially regulated by SlBES1 (Figure 1E). The expression pattern of some of those genes, such as *PG2* and *CSLC12*, varied in different development stages, whereas the others, such as *PE1*, *CD2*, and *XTHs*, were significantly expressed at only one stage. Only the expression levels of *PMEU1* were increased at both B and R stages, suggesting that SlBES1 might affect fruit softening through regulating *PMEU1*, a gene involved in pectin metabolic pathway.

Taken together, these results demonstrated that the BR signaling pathway component SlBES1 positively regulates fruit softening through influencing pectin and other metabolic pathways.

### SlBES1 inhibited PMEU1-associated pectin metabolic pathway

Previous studies have established the role of PME, especially PMEU1, in modulating fruit softening (Tieman and Handa, 1994; Phan et al., 2007). Fruit with low total PME activity shows a reduced degree of methylesterification, decreased fruit firmness, and complete loss of fruit integrity (Tieman et al., 1992; Tieman and Handa, 1994; Phan et al., 2007). Three tomato PME genes, *PME ubiquitously 1* (*PMEU1*), *PE1*, and *PE2*, have been identified and their function in fruit firmness has been proved (Gaffe et al., 1997; Hall et al., 1994). Antisense inhibition of *PMEU1* reduces the total PME activity and accelerates fruit softening (Phan et al., 2007), but *PE1* and *PE2* have no such effect on firmness of tomato fruit (Wen et al., 2013). Our RNA-seq results also indicated that *PMEU1* and *PE1* were up-regulated in firmer *SlBES1-RNAi* fruits (Figure 1E). As discussed above, *SlBES1-OX* fruits phenocopy the fruit with repressed PME activity. Based on these results, we hypothesize that fruit softening in *SlBES1-OX* lines was caused by reduction of PME, mostly PMEU1, which is involved in pectin metabolic pathway

To test this hypothesis, we measured the change in pectin metabolic pathway in *SlBES1* transgenic fruit. The total PME activity was reduced in *SlBES1-OX-3* and *SlBES1-OX-8* from mature green (MG) to R stage, and it was increased in *SlBES1-RNAi-8* and *SlBES1-RNAi-9* from B to R stage (Figure 2A). PME catalyzes the demethylesterification of pectin, and then attenuates the degree of pectin methylesterification (DM) (Senechal et al., 2014; Figure 2C). As the total PME activity decreased, the DM of soluble pectin was increased in *SlBES1-OX-3* and *SlBES1-OX-8* from MG to R stage and decreased in *SlBES1-RNAi-8* and *SlBES1-RNAi-9* from B to R stage (Figure 2B). Additionally, the content and location of highly methylesterified HG, de-esterified HG, and “egg box” were identified through the immunofluorescence of LM20, LM19, and 2F4, respectively (Figure 2D). Compared with the wild type, the signal of methylesterified HG was higher in *SlBES1-OX* and lower in *SlBES1-RNAi* (Figure 2D), whereas the signal of de-esterified HG was lower in *SlBES1-OX* and higher in *SlBES1-RNAi* fruit (Figure 2D), consistent with the total PME activity and DM in *SlBES1* transgenic fruit. The “egg box” signal was significantly elevated in *SlBES1-RNAi* and lowered in *SlBES1-OX* when compared with that in the wild type (Figure 2D). Given the positive role of the “egg box” in cell wall strengthening (Senechal et al., 2014; Silva-Sanzana et al., 2019), the immunolocalization results suggested that decreased content of the “egg box” structure in *SlBES1* fruit was the direct reason for *SlBES1-*induced fruit softening. Calcofluor-white staining showed no major differences in cell size or patterning between transgenic lines (Supplemental Figure S5). The reduced activity of PME and decreased content of its products, as revealed by DM and immunolocalization experiments, support our hypothesis that PME, which is involved in the pectin metabolic pathway, was repressed in *SlBES1-OX* fruit.

**Figure 2.**
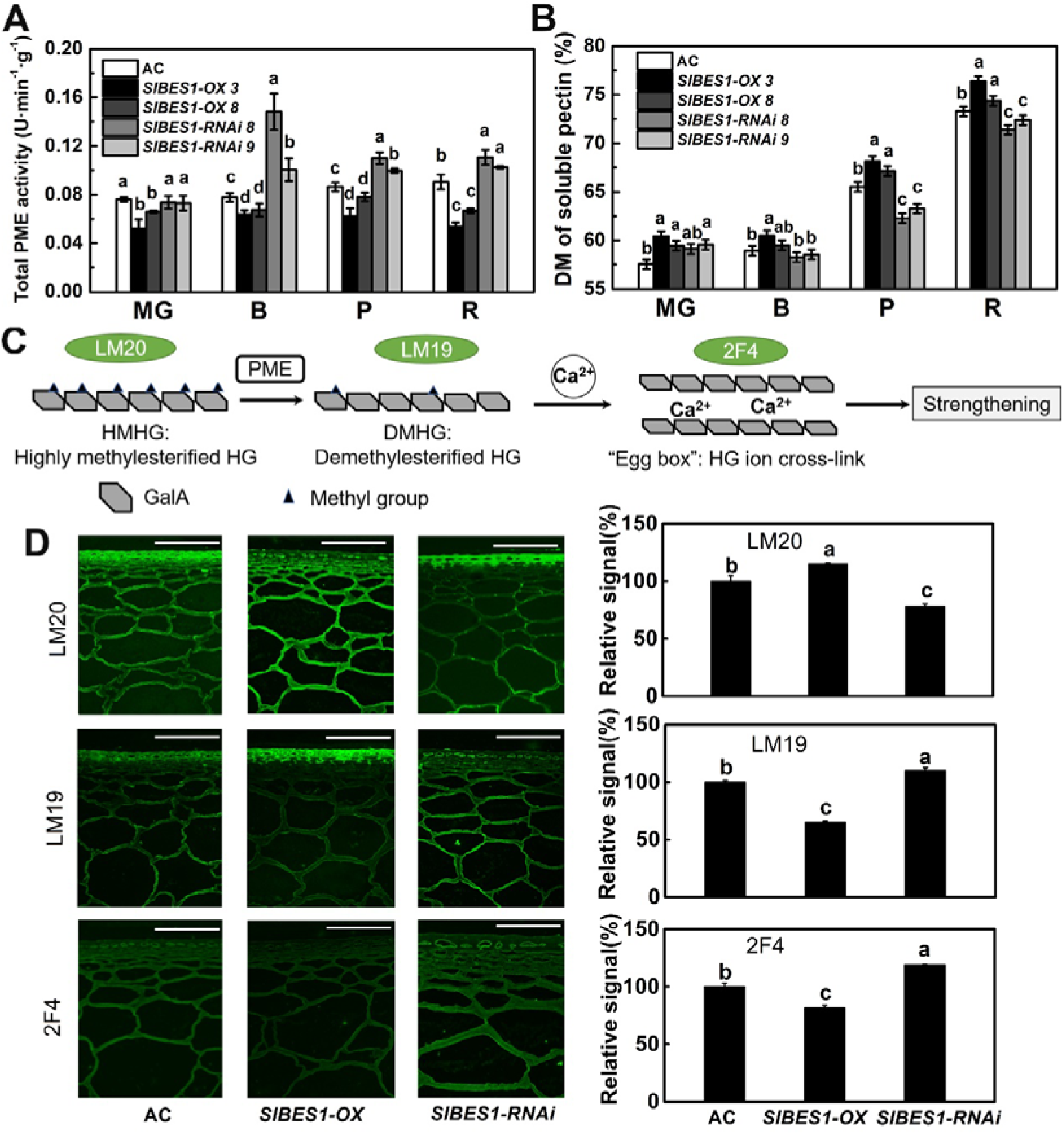
SlBES1 Represses the Pectin Metabolic Pathway. (A-B) Total pectin methylesterase (PME) activity (A) and degree of methylesterification (DM) of soluble pectin (B) in *SlBES1-OX* and *SlBES1-RNAi* fruits at different development stages. MG, mature green stage. B, breaker stage. P, pink stage. R, red ripe stage. (C) Schematic representation of the relationship between pectin methylesterase (PME) and cell wall strengthening. (D) Immunolocalization of highly methylesterified HG (homogalacturonan), de-esterified HG and Ca^2+^ cross-linked HG in *SlBES1-OX* and *SlBES1-RNAi* fruits at red ripe stage. Monoclonal LM20 antibody probe recognizing highly methylesterified HG, LM19 probe recognizing de-esterified pectin, and 2F4 probe recognizing Ca^2+^ cross-linked HG were used to label tomato pericarp tissue. Representative sections of fruits from each of AC (wild type), *SlBES1-OX-3* and *SlBES1-RNAi-8* lines are presented. Scale bar represents 100 μm. The right graphs show the relative fluorescence signal of each antibody. Relative signals are calculated through software ImageJ. Values were normalized with respect to AC (wild type). In (A-B) and (D) Each data point represents means±SD of three determinations. Different letters indicate significant difference between different groups (one-way ANOVA with Tukey’s test, *p*<0.05).

Then, the PME gene required for the fruit softening process was investigated. To date, *PMEU1*, has been proved to be the only PME gene that affects both total PME activity and fruit firmness in tomato (Phan et al., 2007). The expression level of *PMEU1* was decreased in *SlBES1-OX* from the MG to the R stage and increased in *SlBES1-RNAi* from the B to the R stage (Figure 3A); this corresponds to the total PME activity and DM changes (Figure 2A and B). Interestingly, the expression level of *PMEU1* was also down-regulated in EBL-treated fruits, while EBL treatment offset the down-regulated *PMEU1* expression in *d^im^* (Supplemental Figure S6A and B), suggesting a role of BR in repressing *PMEU1* expression. These results indicated that both *PMEU1* mediated pectin metabolic pathway and the expression level of *PMEU1* was repressed by SlBES1. All genetic and molecular results suggest that SlBES1 represses the expression of *PMEU1* to promote fruit softening.

**Figure 3.**
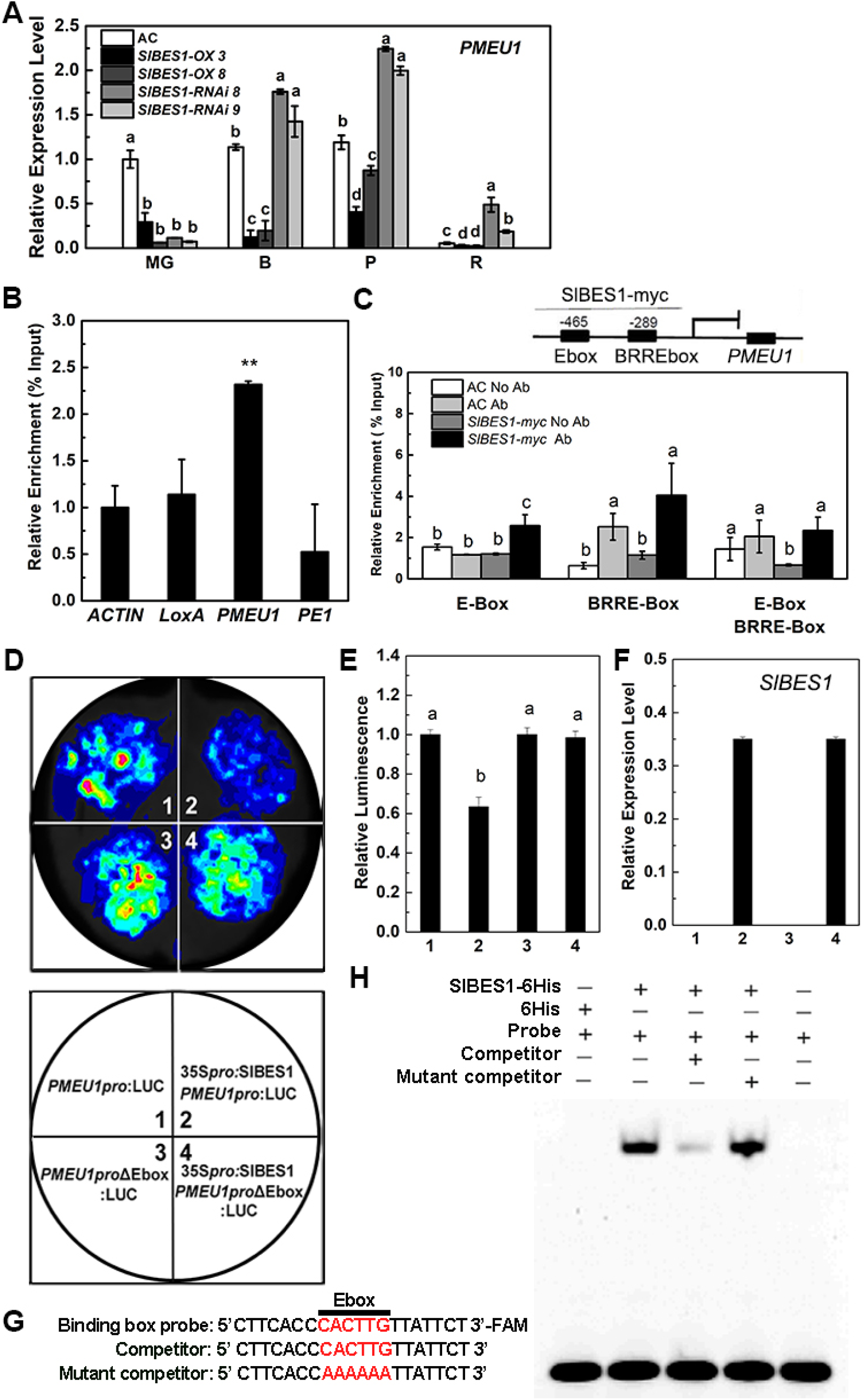
SlBES1 Represses the Transcriptional Expression of *PMEU1*. (A) Relative expression levels of *PMEU1* in *SlBES1-OX* and *SlBES1-RNAi* fruits at different development stages. MG, mature green stage. B, breaker stage. P, pink stage. R, red ripe stage. Each data point represents means±SD of three determinations. Different letters indicate significant difference between different groups (one-way ANOVA with Tukey’s test, *p*<0.05). (B) ChIP-qPCR assays showing that SlBES1 associated with the locus of candidate genes, *LoxA*, *PMEU1* and *PE1*. Chromatin of transgenic plants expressing *35Spro:SlBES1-myc* (*SlBES1-myc*) was immunoprecipitated with anti-myc antibody, and results of gene *ACTIN7* served as control. The relative enrichment for ChIP signal was displayed as the percentage of total input DNA. *PMEU1* locus contains both E-box and BRRE-box. Values are means ± SD of three biological replicates. Statistical analysis was performed with ANOVA. Bars with asterisks indicates significant difference (\**p*<0.05). (C) The upper graph, schematic diagram of *PMEU1*, indicating the amplicons used for ChIP-qPCR. Position of E-box and BRRE-box (BR Response Element) are indicated. The below graph, ChIP-qPCR assays, showing that SlBES1 associated with *PMEU1* locus in tomato fruits. The relative enrichment for SlBES1 at two *PMEU1* promoter motifs, E-box and BRRE-box, was calculated against the *ACTIN2* promoter. AC (wild type) without and with antibody were set as blank control and negative control, respectively. *SlBES1-myc* without antibody was also set as negative control. The mean value of two technical replicates was recorded for each biological replicate. Values are means ±SD of three biological replicates. Different letter indicates a significant difference among groups for each locus (one-way ANOVA with Tukey’s test, *p*<0.05). (D) Transient expression assays showing that SlBES1 represses *PMEU1* expression. Representative images of *N. benthamiana* leaves are taken 48 h after infiltration. The bottom panel indicates the infiltrated constructs. (E-F) Luminescence intensity (E) and *SlBES1* expression level (F) in different treatment as indicated in (D). Values are means ± SD of six biological replicates. Different letter indicates a significant difference among groups for each locus (one-way ANOVA with Tukey’s test, *p*<0.05). (G) Motif sequence of labeled, unlabeled competitor and mutant competitor probes. Mutant competitor in which the 5’-CACTTG-3’ motif was replaced with 5’-AAAAAA-3’. (H) DNA electrophoretic mobility shift assay (EMSA) showing that the SlBES1-His binds to E-box of *PMEU1* in *vitro*. FAM-labeled probes were incubated with SlBES1-His and the free and bound DNAs were separated on an acrylamide gel.

### SlBES1 mediated direct transcriptional inhibition of *PMEU1* to confers fruit softening

To test the hypothesis of possible inhibited expression of *PMEU1* by SlBES1, we carried out further experiments to prove the transcriptional regulation of SlBES1 in the expression of *PMEU1*. We first selected several candidate genes including *PMEU1* on the basis of our results and previous proteomics data (Liu et al., 2016). ChIP-qPCR results showed a significantly higher enrichment for the promoters of *PMEU1* than for *PE1* (Figure 3B). Transcriptional factor BES1 preferentially binds to E-box (CANNTG) or BRRE-box (CGTG(T/C)G) to regulate the expression of downstream genes in *Arabidopsis* (Yu et al., 2011). The two binding sites, E-box and BRRE-box, were located at −465 bp and −289 bp of the *PMEU1* promoter as predicted by Jaspar database (http://jaspar.genereg.net/) (Figure 3C). The ChIP-qPCR results further indicated enrichment of SlBES1 binding to the region containing E-box but not to that of the BRRE-box (Figure 3C), suggesting that SlBES1 might regulate *PMEU1* expression by binding to the E-box.

To test if E-box is necessary for SlBES1 regulation of the *PMEU1* gene, we generated a *PMEU1pro:LUC* reporter, in which luciferase (LUC) was fused with the PMEU1 promoter. Co-expression of *PMEU1pro:LUC* with the *35Spro:SlBES1* construct led to a significantly reduced luminescence intensity, suggesting that SlBES1 represses the expression of *PMEU1*. Importantly, when *PMEU1pro:LUC* was replaced by the *PMEU1proΔEbox:LUC*, in which the E-box of the *PMEU1* promoter involved in the transcriptional regulation of SlBES1 on *PMEU1* was deleted, SlBES1-mediated repression of the *PMEU1* expression was abolished (Figure 3D to F).

We then conducted electrophoretic mobility shift assay (EMSA) with purified SlBES1-His and a 20-bp DNA probe containing this E-box motif. SlBES1-His bound to the DNA probe, and this binding was successfully outcompeted by unlabeled DNA probe but not by the DNA probe without E-box. These results suggested that SlBES1 directly binds to the *PMEU1* promoter through E-box (Figure 3G and H). E-box was the binding site of SlBES1 to repress gene expression in tomato. Although BES1/BZR1 mostly bind to E-box to activate gene expression and to BRRE to repress genes, there are exceptions (Nolan et al., 2020). A recent study also showed that OsBZR1 bind to E-box to inhibit downstream gene expression in rice (Fang et al., 2020).

Silencing *PMEU1* caused softer fruits with lower total PME activity and exhibited a complete loss of fruit integrity and shorter shelf life (Figure 4A to D), which was in accordance with previous studies (Tieman and Handa, 1994; Phan *et al*., 2007). More importantly, knocking down *PMEU1* in the *SlBES1-RNAi* background (TRV-*PMEU1/SlBES1-RNAi*) suppressed the *SlBES1-RNAi* phenotypes of higher firmness and the longer shelf life. Fruit firmness, shelf life, and total PME activity of TRV-*PMEU1/SlBES1-RNAi* were almost the same as wild type, indicating that SlBES1-mediated fruit softening is mainly mediated by the inhibition of *PMEU1* (Figure 4 A to D).

**Figure 4.**
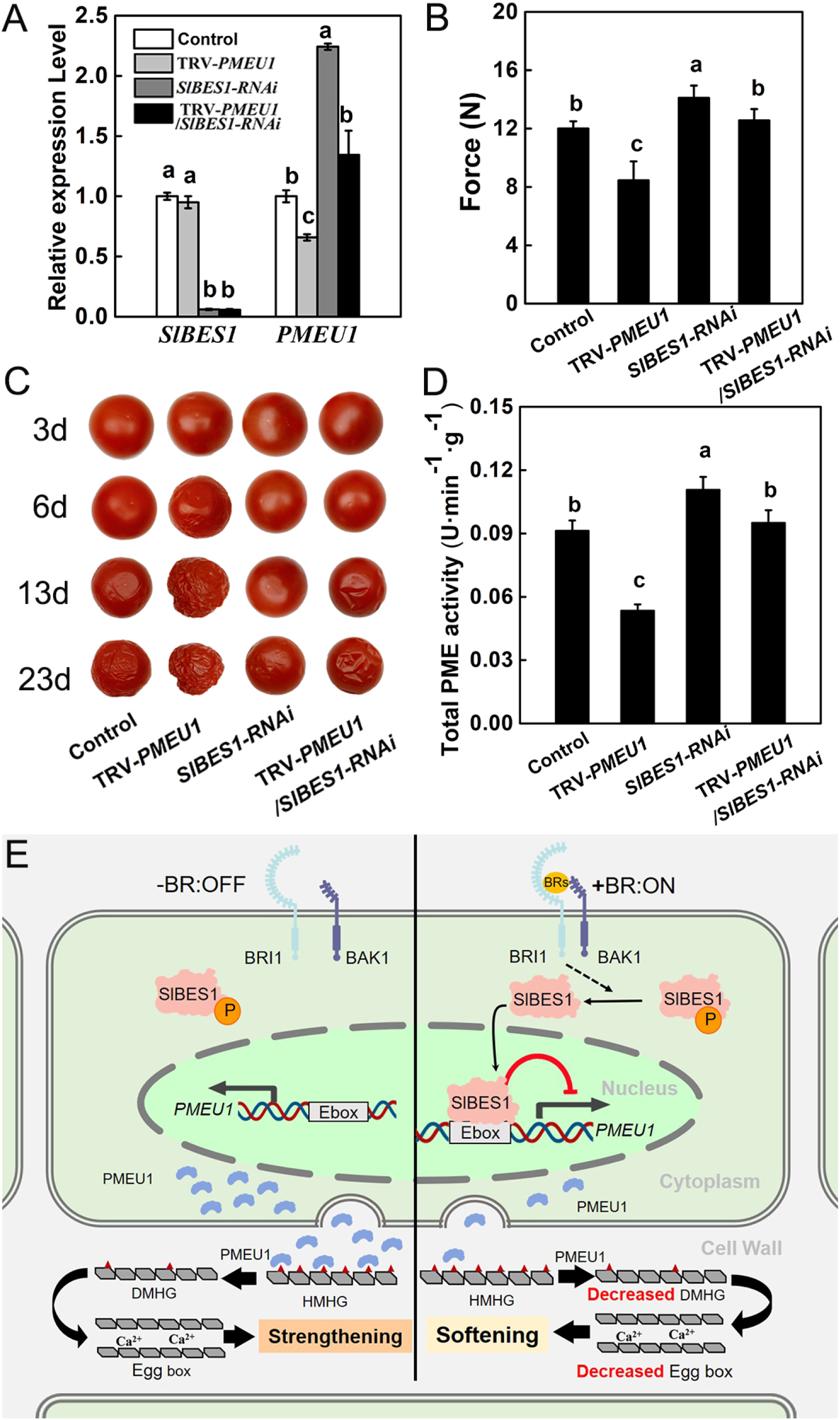
Effects of Silencing *PMEU1* in AC and *SlBES1-RNAi* on Tomato Firmness and Shelf Life. (A) Relative expression levels of *SlBES1* and *PMEU1* in fruits of AC (Control), virus-induced *PMEU1* silencing lines (TRV-*PMEU1*), *SlBES1-RNAi*, and virus-induced *PMEU1* silencing in *SlBES1-RNAi* (TRV-*PMEU1*/*SlBES1-RNAi*) at red ripe stage. (B) Fruit firmness of Control, TRV-*PMEU1*, *SlBES1-RNAi*, and TRV-*PMEU1*/*SlBES1-RNAi* at red ripe stage. Values are means ± SD of twenty-four biological replicates. (C) The tomato shelf life of Control, TRV-*PMEU1*, *SlBES1-RNAi*, and TRV-*PMEU1*/*SlBES1-RNAi*. Fruits were harvested at red ripe stage and stored at room temperature. The progression of fruit deterioration was recorded by time-lapse photography. Time after harvest is specified by days. The storage condition was 25◻ and 35% relative humidity. (D) Total PME activity of Control, TRV-*PMEU1*, *SlBES1-RNAi*, and TRV-*PMEU1*/*SlBES1-RNAi* fruits at red ripe stage. (A) to (D) Expected for fruit firmness, values are means ± SD of three biological replicates. Different letter indicates a significant difference among groups (one-way ANOVA, *p*<0.05, Tukey’s test). (E) Schematic representation of relationship between transcriptional regulation of slbes1 on *pmeu1* and fruit firmness. When BRs are absent as left showed, receptor BRI and co-receptor BKI1 are separated. Transcriptional factor SlBES1 with phosphate group couldn’t inhibit pectin methylesterase gene *PMEU1* expression. Enzyme protein PMEU1 is formed and secreted into the cell wall. Then PMEU1 de-esterified HMHG and produces DMHG, DMHG formed “Egg box” through cross-link with Ca^2+^. Egg box provides strengthening support to fruit firmness. When BRs are present, BRs bind to BRI and BKI1. A series of phosphorylation and dephosphorylation are occurred, which lead to the dephosphorylation of SlBES1. Dephosphorylated SlBES1 binds to the E-box of *PMEU1* promoter and then represses the expression of *PMEU1*. Fewer PMEU1 are secreted into the cell wall and pectin methylesterase activity is attenuated, so the contents of DMHG and Egg box are decreased which causing fruit softening. BRI1, BRASSINOSTEROID INSENSITIVE 1; BAK1, BRI1-ASSOCIATED KINASE1; BES1, BRI1-EMS-SUPPRESSOR 1; PMEU1, PECTIN METHYLESTERASE UBIQUITOUSLY 1; HMHG, highly methylesterified HG; DMHG, demethylesterified HG.

Taken together, these results demonstrated that SlBES1 directly binds to the E-box of *PMEU1* to repress its expression, thereby promoting fruit softening.

### Gene editing of *SlBES1* by CRISPR/Cas9 enhances fruit firmness without negative effect on nutritional quality

Above results suggested *SlBES1* may be a potential gene that can be used for breeding tomatoes with longer shelf life and maintaining optimum flavor. Then we generated *SlBES1-KO* by using CRISPR/Cas9 through targeting the 5’-TAGTTGGTGATGAAAGAGG TGG-3’ in the second extrinsic of SlBES1 antisense strand with *pYLCRISPR/Cas9* (Ma et al., 2015), and the result showed that the fruit of *SlBES1-KO*, a *SlBES1* knockout mutant line, had higher firmness and longer shelf life without loss of nutritional quality (Supplemental Figure S7 and Table 3).

In summary, this study provides genetic, biochemical, and molecular evidence for SlBES1-mediated repression of the pectin metabolic pathway and fruit softening. At the transcriptional level, SlBES1 directly binds to the E-box of the *PMEU1* promoter to repress *PMEU1* expression, thereby promoting fruit softening (Figure 4 E). BR also plays a positive role in this process. Fruits with silenced *SlBES1* described in this study provide a promising solution to improve fruit firmness and extended shelf life without negative effect on visual and nutritional quality.

## Methods

### Plant Materials and Growth Conditions

All transgenic lines were constructed in tomato cultivars Ailsa Craig (AC). Cultivars *Lycopersicon pimpinellifolium* (Spim), Condine Red (CR), and Craigella are the parental line of *cu-3* (Scheer et al., 2003), *d^im^*, and *d*^x^ (Li et al., 2015), respectively. Plants were cultivated under a 16 h photoperiod (22/28 ◻, night/day). Fruits ripening stages were defined as described previously (Giovannoni, 2004). More details are described in Supplemental Methods.

### Vector Constructs and Plant Transformation

Recombinant plasmid *35S pro:SlBES1-myc* and *pBIN19-SlBES1-RNAi* were used for *SlBES1-OX* and *SlBES1-RNAi* construct, respectively. Constructs were introduced into AC via *Agrobacterium*-mediated transformation (Shao et al., 2019). Homozygous T2 transgenic plants were selected based on resistance to kanamycin and then used for further tests. More details are described in Supplemental Methods.

### Generation of the CRISPR/Cas9 Mutant

The guide RNA sequence CCACCTCTTTCATCACCAACTA for *SlBES1* was cloned into *pYLCRISPR/Cas9* as described previously (Ma et al., 2015). After inducing construct into AC by *Agrobacterium*-mediated transformation, sequences containing guide RNA target sites were amplified and sequenced to confirm mutations in targeted regions. Homozygous T2 transgenic plants were selected for further experiments.

### Chemical Treatment

AC and *d^im^* at mature green stage were treated with 24-epibrassinolide (EBL, Sigma, St. Louis, MO) as described previously (Liu et al., 2014) and then collected at 1st, 3rd, 6th, and 9th day for further tests. Tomato seeds were sown in half-strength MS containing 0.5 μM propiconazole (Pcz, Sigma, St. Louis, MO). Plants were grown at 28 ◻ with total darkness for 6 d, then the lengths of seedlings hypocotyls and roots were measured through software ImageJ after photographed.

### Firmness Determination

Firmness was tested at the fruit equatorial region with a texture analyzer (TA-XT2i, Godalming, UK) and a 7.5 mm probe with 1 mms^−1^ penetration speed as described previously (Liu et al., 2018).

### RNA Extraction and Relative Quantitative PCR

For RNA extraction, 0.1g of leaves or fruits was mixed with 1 mL RNA iso plus according to manufacturer’s instruction (Takara, Kusatsu, Japan), then RNA was reverse-transcribed into cDNA using PrimeScript RT reagent with gDNA Eraser (Takara, Kusatsu, Japan). TB Green (Takara, Kusatsu, Japan) was then used in Step One Real-Time PCR System (Applied biosystem, CA, USA) for relative quantitative PCR (qPCR). The gene-specific primers used were listed in Supplemental Table 2.

### RNA-Seq

AC and *SlBES1-RNAi-8* fruits at breaker and red ripe stages were collected for total RNA extraction and Illumina MiSeq library was constructed as described by manufacturer’s instructions (Illumina, San Diego, CA, USA) and then sequenced with the Illumina Miseq platform.

### ChIP-qPCR, Transient Expression Assay in Tobacco and EMSA

Chromatin immunoprecipitation quantitative PCR (ChIP-qPCR) for fruits was performed following previous reports (Liu et al., 2019). Used primer pairs were listed in Supplemental Table 2. Transient expression assay in tobacco (*Nicotiana benthamiana*) and EMSA was performed as described previously (Shao et al., 2019). More details are described in Supplemental Methods.

### Determination of PME Enzyme Activity and Degree of Pectin Methylesterification

Total PME activity and degree of pectin methylesterifcation (DM) were determined as previously reported (Tucker et al., 1992; Freitas et al., 2012; Kyomugasho et al., 2015; Chylinska et al., 2016). More details are described in Supplemental Methods.

### Immunofluorescence

Immunofluorescence was performed as described previously (Silva-Sanzana et al., 2019). Fresh pericarps were fixed with FAA and submerged with LR White resin (Sigma, St. Louis, MO) to obtain sections for treating with LM20, LM19, and 2F4 antibodies (Plantprobes, Leeds, UK). More details are described in Supplemental Methods.

### Virus-Inducing Gene Silencing

A coding region fragment of *PMEU1* (1-300 bp) was inserted into pTRV2 to generate pTRV2-*PMEU1*. pTRV1 and pTRV2-*PMEU1* were induced into AC fruits following the previous protocol (Fu et al., 2005).

### Carotenoid and Ascorbic Acid Analysis

The concentrations of carotenoid and ascorbic acid were determined as described previously (Liu et al., 2018).

### Data Availability

The RNA-seq data of this study were submitted to the NCBI Sequence Read Archive (SRA) database (http://www.ncbi.nlm.nih.gov/sra/) with the BioProject ID PRJNA635540.

## ACCESSION NUMBERS

The accession numbers for the genes described in this report are as follows: *SlBES1* (Solyc04g079980), *PMEU1* (Solyc03g123630), *PE1* (Solyc07g064170), *SlDWARF* (Solyc02g089160), *SlCPD* (Solyc06g051750), *SlCYP724B2* (Solyc07g056160), *SlCYP90B3* (Solyc02g085360), *LOXA* (Solyc08g014000), *ACTIN2* (Solyc11g005330).

## FUNDING

This research was supported by National Natural Science Foundation of China (Key Program, No.31830078), China Postdoctoral Science Foundation (No. 2019M662067) and Zhejiang Provincial Ten-thousand Program for Leading Talents of Science And Technology Innovation (2018R52026).

## AUTHOR CONTRIBUTION

Q.W., C.L., and H.L. designed the research. H.L., M.Z., C.J., M.Q., S.H., F.M., Z.S., and D.L. performed the research. H.L. D.L., and Y.L. analyzed data. H.L., L.L, C.J., and Q.W. wrote the manuscript.

## ACKNOWLEDGEMENTS

We thank Tomato Genetics Resource Center (University of California, Davis, CA) for providing tomato seeds used in our research. We also thank Prof. Daqi Fu (China Agricultural University) and Prof. Yaoguang Liu (South China Agriculture University) for kindly providing vectors for VIGS and pYLCRISPR/Cas9 plasmids. No conflict of interest declared.

## Notes

### Competing Interest Statement

The authors have declared no competing interest.

