## supplemental figure S1-S7 for "Brassinosteroid signaling component SlBES1 promotes tomato fruit softening through transcriptional repression of *PMEU1*"

**Brassinosteroids signaling component SlBES1 targeted promotes tomato fruits softening through transcriptional repression on *PMEU1***

Haoran Liu^1+^, Lihong Liu^1+^, Dongyi Liang^1^, Min Zhang^1^, Chengguo Jia^3^, Mingfang Qi^4^, Yuanyuan Liu^1^, Zhiyong Shao^1^, Fanliang Meng^1^, Songshen Hu^1^, Chuanyou Li^2^*, Qiaomei Wang^1^*.

1 Key Laboratory of Horticultural Plant Growth and Development, Ministry of Agriculture, Department of Horticulture, Zhejiang University, Hangzhou 310058, China

2 State Key Laboratory of Plant Genomics, National Centre for Plant Gene Research (Beijing), Institute of Genetics and Developmental Biology, Chinese Academy of Sciences, Beijing 100097, China

3 College of Plant Science, Jilin University, Changchun 130062, Jilin, China

4 Key Laboratory of Protected Horticulture of Ministry of Education, College of Horticulture, Shenyang Agricultural University, Shenyang 110866, China

^+^These authors contributed equally to this article.

**Supplementary Figure Legends**

**Supplementary Figure S1-S7**

**Supplemental Methods**

**Supplementary Figure Legends**

**Figure S1. Tomato gene, *SlBES1*, is homologous gene of *AtBES1*.**

(A) Phylogenetic tree of tomato and *Arabidopsis* BES1 proteins. The phylogenetic tree is constructed based on the complete protein sequence alignment by the Neighbor-Joining method with bootstrapping analysis (1000 replicates) using MEGA X software.

(B) Protein alignment of tomato gene *SlBES1* and *Arabidopsis* gene *AtBES1*. Nuclear localization signal domain, BIN2 catalyzing site and PEST domain were indicated with black lines.

(C) Expression analysis of *SlBES1* and *SlDWARF* (BR biosynthetic gene) in *d^x^*, a mutant impaired in *SlDWARF*, and its wild type (Craigella) as well as BR receptor BRI1 mutant *cu-3* and its wild type, Spim. Data shown represents means ±SD of three biological replicates. Asterisks indicate significant difference compared with wild-type (Student’s *t*-test, *p* < 0.05).

**Figure S2 SlBES1 has conserved functions of BES1.**

(A) Phenotypes of AC (wild type), *SlBES1-OX* and *SlBES1-RNAi* seedlings grown on half-strength MS medium with or without 0.5 μM Pcz in the dark, photographs were taken 6 days after sowing. Scale bar = 1 cm.

(B) Hypocotyl length and root length of AC (wild type), *SlBES1-OX* and *SlBES1-RNAi* seedlings grown on half-strength MS medium with or without Pcz in the dark for 6 days. Data shown are means ±SD of three biological replicates. Asterisks indicate significant difference compared with wild type (one-way ANOVA, *p*<0.05, Tukey’s test).

(C) Relative gene expression levels of BR biosynthetic genes, *SlDWARF*, *SlCPD*, *SlCYP724B2*, *SlCYP90B3* in fruits of *SlBES1-OX* and *SlBES1-RNAi*. L, leaves. MG, mature green. B, breaker. P, pink. R, red ripe. Data shown represents means ±SD of three biological replicates. Asterisks indicate significant difference compared with wild type (one-way ANOVA, *p*<0.05, Tukey’s test).

**Figure S3. Ethylene production of *SlBES1-OX* and *SlBES1-RNAi* tomato fruits at different development stages.**

MG, mature green. B, breaker. P, pink. R, red ripe. Data shown represents the means ±SD of three biological replicates from at least six independent fruits. Different letters indicate significant difference compared to AC (wild type) (one-way ANOVA with Tukey’s test, p<0.05).

**Figure S4. Fruit firmness of fruits at different days after 2,4-epibrassinolide (EBL) treatment.**

Fruits firmness of AC fruits (A) as well as BR biosynthetic deficient mutant *d^im^* and its wild type Condine Red (CR) (B) at different days after exogenous treatment of 24-epibrassinolide (EBL). Values are means ± SD of twenty-four biological replicates from at least six independent fruits. Different letter indicates a significant difference among groups (one-way ANOVA, *p*<0.05, Tukey’s test).

**Figure S5. Calcofluor white staining of pericarp sections from *SlBES1-OX* and** ***SlBES1-RNAi*.** Scale bar = 100 μm

**Figure S6. Relative expression levels of *PMEU1* in fruits of AC (A) as well as BR biosynthetic deficient mutant *d^im^* and CR (B) at different days after EBL treatment.** Data shown represents means ±SD of three biological replicates. Asterisks indicate significant difference compared with wild-type (Student’s *t*-test, **p* < 0.05, ** *p* < 0.01).

**Figure S7. Effect of *SlBES1* CRISPR/Cas9 mutant on fruit firmness and shelf life.**

(A) Target sequence that contains mutation of *SlBES1-KO* compared to AC (Wild type).

(B) Fruits firmness of *SlBES1-KO* at red ripe stage (Student’s *t*-test, **p* < 0.05).

(C) AC (Wild type) and *SlBES1-KO* fruits were harvested at pink stage and stored at room temperature. The progression of fruit deterioration was recorded by time-lapse photography. Time after harvest is specified by days.

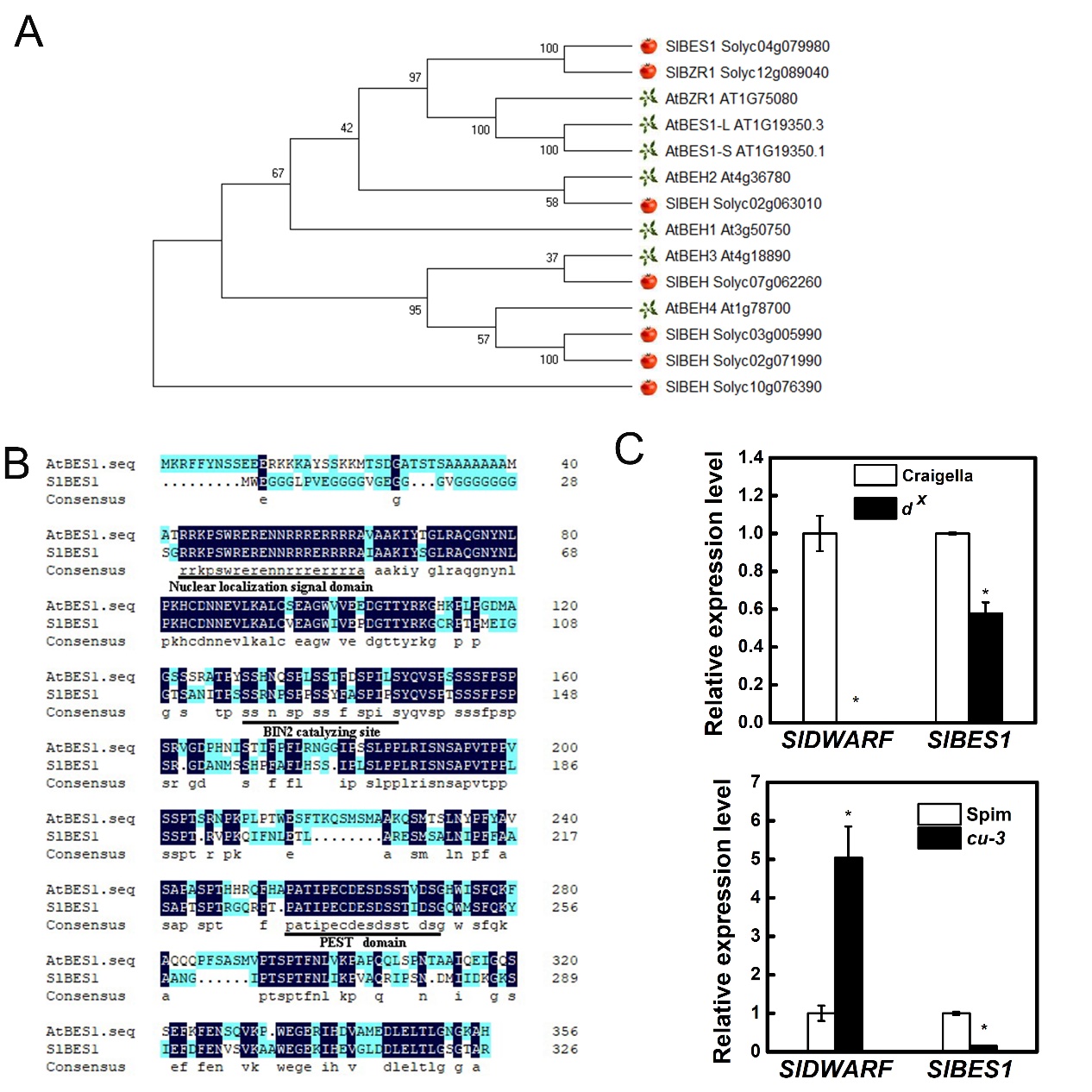

Figure S1. Tomato gene, *SlBES1*, is homologous gene of *AtBES1*.

(A) Phylogenetic tree of tomato and *Arabidopsis* BES1 proteins. The phylogenetic tree is constructed based on the complete protein sequence alignment by the Neighbor-Joining method with bootstrapping analysis (1000 replicates) using MEGA X software.

(B) Protein alignment of tomato gene *SlBES1* and *Arabidopsis* gene *AtBES1*. Nuclear localization signal domain, BIN2 catalyzing site and PEST domain were indicated with black lines.

(C) Expression analysis of *SlBES1* and *SlDWARF* (BR biosynthetic gene) in *d^x^*, a mutant impaired in *SlDWARF*, and its wild type (Craigella) as well as BR receptor BRI1 mutant *cu-3* and its wild type, Spim. Data shown represents means ±SD of three biological replicates. Asterisks indicate significant difference compared with wild-type (Student’s *t*-test, *p* < 0.05).

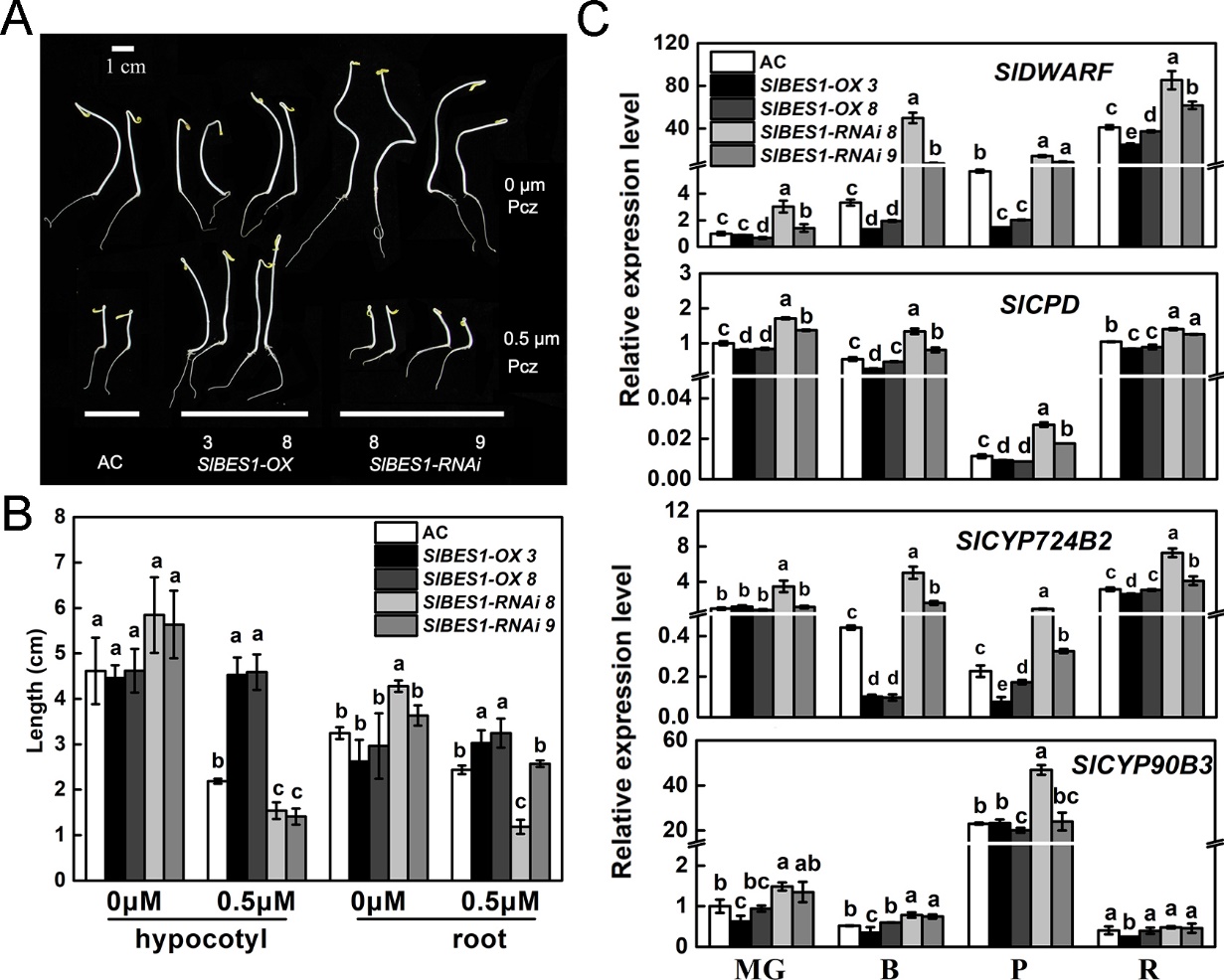

Figure S2 SlBES1 has conserved functions of BES1.

(A) Phenotypes of AC (wild type), *SlBES1-OX* and *SlBES1-RNAi* seedlings grown on half-strength MS medium with or without 0.5μM Pcz in the dark, photographs were taken 6 days after sowing. Scale bar = 1 cm.

(B) Hypocotyl length and root length of AC (wild type), *SlBES1-OX* and *SlBES1-RNAi* seedlings grown on half-strength MS medium with or without Pcz in the dark for 6 days. Data shown are means ±SD of three biological replicates. Asterisks indicate significant difference compared with wild type (one-way ANOVA, *p*<0.05, Tukey’s test).

(C) Relative gene expression levels of BR biosynthetic genes, *SlDWARF*, *SlCPD*, *SlCYP724B2*, *SlCYP90B3* in fruits of *SlBES1-OX* and *SlBES1-RNAi*. L, leaves. MG, mature green stage. B, breaker stage. P, pink stage. R, red ripe stage. Data shown represents means ±SD of three biological replicates. Asterisks indicate significant difference compared with wild type (one-way ANOVA, *p*<0.05, Tukey’s test).

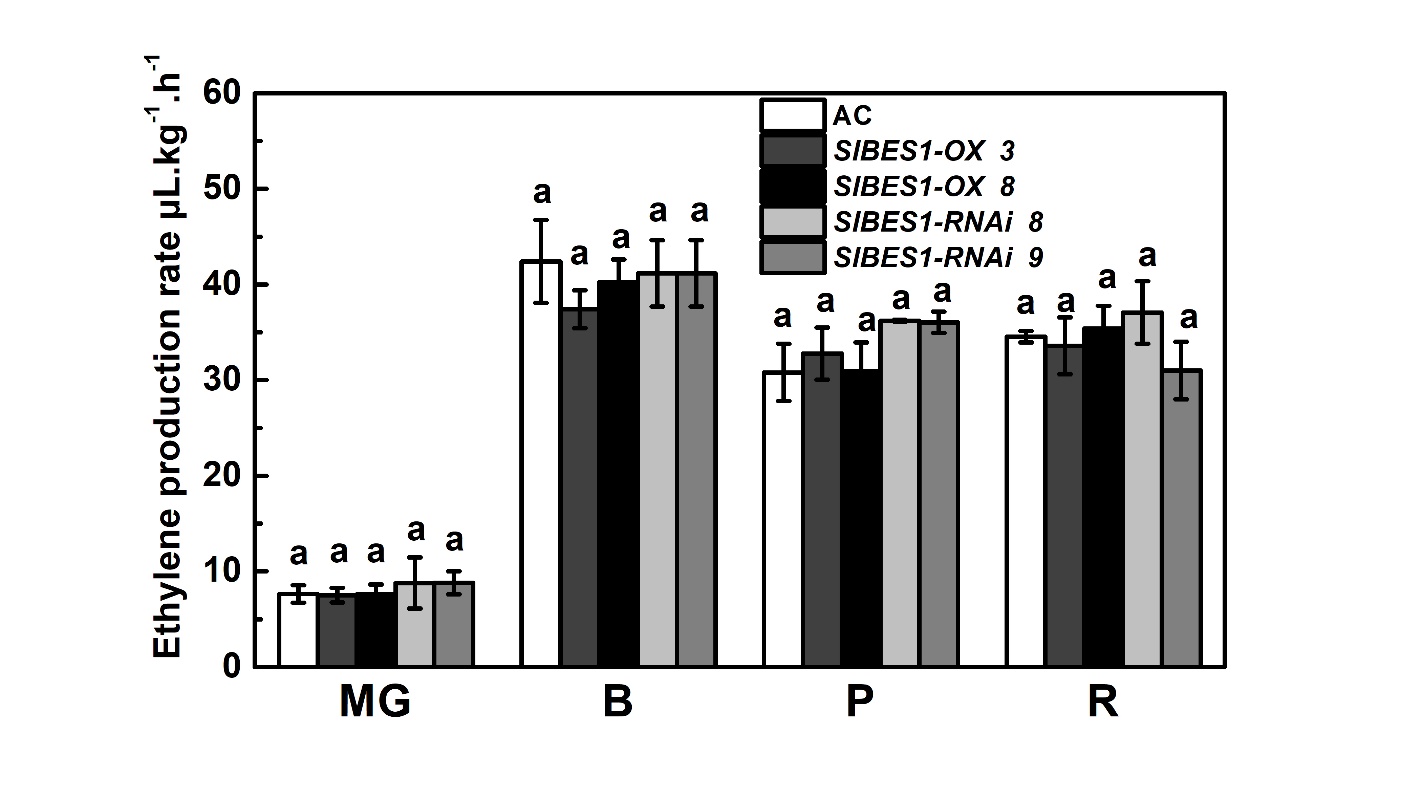

Figure S3. Ethylene production of *SlBES1-OX* and *SlBES1-RNAi* tomato fruits at different development stages. MG, mature green stage. B, breaker stage. P, pink stage. R, red ripe stage. Data shown represents the means±SD of three biological replicates from at least six independent fruits. Different letters indicate significant difference compared to AC (Wild type) (one-way ANOVA with Tukey’s test, p<0.05).

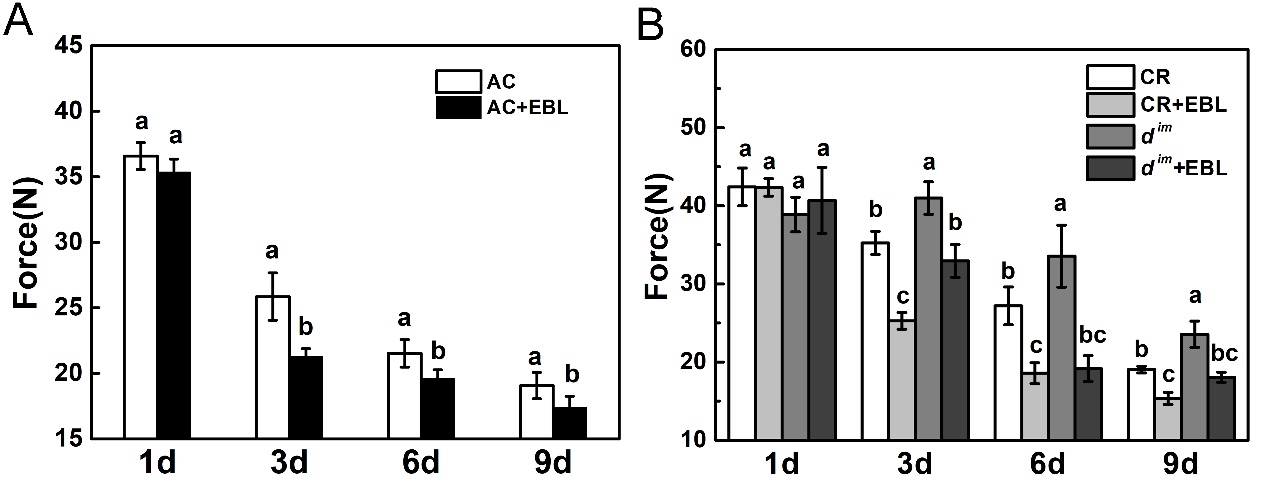

Figure S4. Fruit firmness of fruits at different days after 24-epibrassinolide (EBL) treatment.

Fruits firmness of AC fruits (A) as well as BR biosynthetic deficient mutant *d^im^* and its wild type Condine Red (CR) (B) at different days after exogenous treatment of 24-epibrassinolide (EBL). Values are means ±SD of twenty-four biological replicates from at least six independent fruits. Different letter indicates a significant difference among groups (one-way ANOVA, *p*<0.05, Tukey’s test).

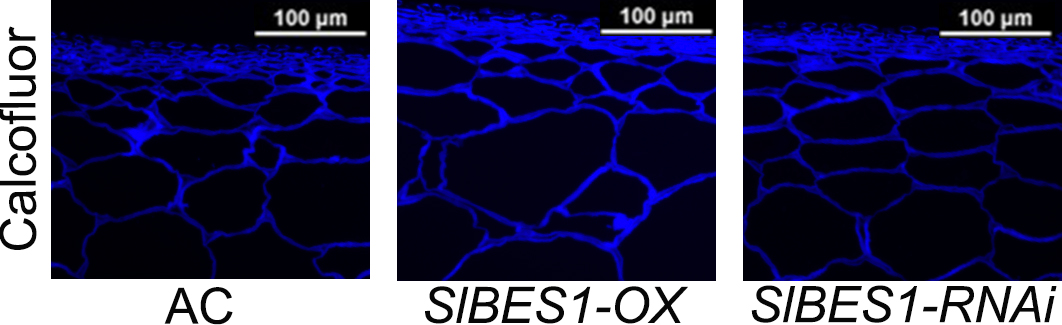

Figure S5. Calcofluor white staining of pericarp sections from *SlBES1-OX* and *SlBES1-RNAi*. Scale bar = 100 μm

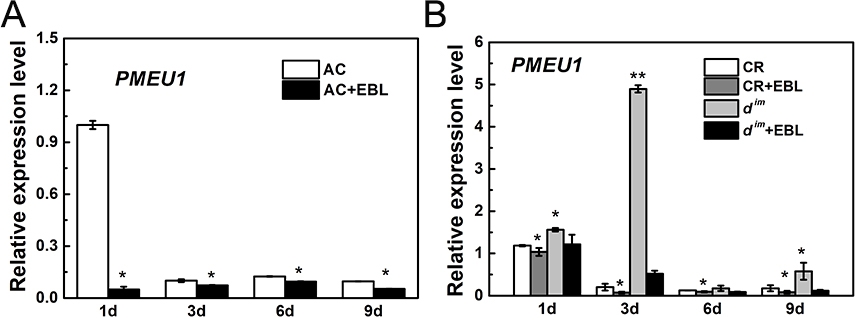

Figure S6. Relative expression levels of *PMEU1* in fruits of AC (A) as well as BR biosynthetic deficient mutant *d^im^* and CR (B) at different days after EBL treatment. Data shown represents means ±SD of three biological replicates. Asterisks indicate significant difference compared with wild-type (Student’s *t*-test, **p* < 0.05, ** *p* < 0.01).

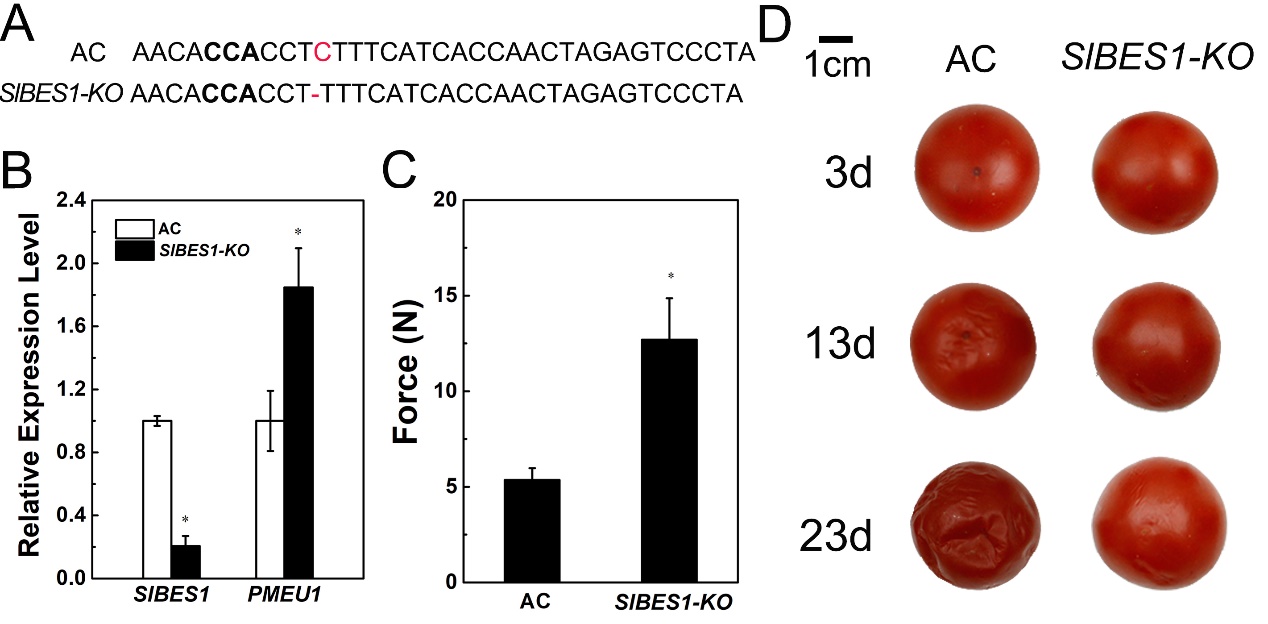

Figure S7. Effect of SlBES1 CRISPR/Cas9 mutant on fruit firmness and shelf life.

(A) Target sequence that contains mutation of *SlBES1-KO* compared to AC (Wild type).

(B) Relative expression levels of *SlBES1* and *PMEU1* in fruits of *SlBES1-KO* (Student’s *t*-test, **p* < 0.05).

(C) Fruits firmness of *SlBES1-KO* at red ripe stage (Student’s *t*-test, **p* < 0.05).

(D) AC (Wild type) and *SlBES1-KO* fruits were harvested at pink stage and stored at room temperature. The progression of fruit deterioration was recorded by time-lapse photography. Time after harvest is specified by days.

**Supplemental Methods**

**Plant Materials and Growth Conditions**

All transgenic lines were constructed in tomato (*Solanum lycopersicum*) cultivars Ailsa Craig (AC). Cultivars Condine Red (CR) and Craigella are the parental line of BR biosynthesis deficiency mutant *d^im^* and *d*^x^, respectively (Li et al., 2015). Lycopersicon pimpinellifolium (Spim) is the wild type of BR receptor mutant, *cu-3* (Scheer et al., 2003). Above tomato plants were cultivated in a greenhouse under a 16 h photoperiod (22/28 ℃, night/day). The number of tagged flowers at anthesis was limited to fewer than five per cluster. The ripening stages, including mature green stage (MG), breaker stage (B), pink stage (P), and red ripe stage (R), of fruits were defined based on fruit color as described previously (Giovannoni, 2004). After firmness determination and immunofluorescence**,** three biological replicates for transgenic fruits pieces (each biological replicate consisting of three fruits) or fully expanded leaves from 4-week-old BR mutants were frozen with liquid nitrogen as samples, and then stored at -80℃ for further tests.

**Vector Constructs and Plant Transformation**

Vector constructs for transgenic were generated following standard molecular biology protocols. For *SlBES1-OX* plants, the full-length coding sequence of *SlBES1* without termination codon was amplified via PCR and inserted into the pGWB17 vector using Gateway (Invitrogen) technology (Nakagawa et al., 2007) to get the *Pro35S:SlBES1-myc* construct. For *SlBES1-RNAi* construct, a fragment of *SlBES1* with length of 305bp was amplified and then inserted into intermediate vector pHANNIBAL in the positive orientation, thereby generating vector *pHANNIBAL-SlBES1*. The same fragment of *SlBES1* was inserted into *pHANNIBAL-SlBES1* in the reverse orientation, generating the vector *pHANNIBAL-SlBES1i*. The target inverted repeat sequences were obtained by *Sac*I and *Spe*I digestion of *pHANNIBAL-SlBES1i*. Eventually, these sequences were inserted into *pBIN19* under the control of CaMV 35S promoter to generate *pBIN19-SlBES1-RNAi*.

The above constructs were introduced into tomato cultivars AC via *Agrobacterium tumefaciens* LBA4404-mediated transformation (Shao et al., 2019). According to screening for regenerated shoots, T1 transgenic plants were then selected based on their resistance to Kanamycin. Homozygous T2 or T3 transgenic plants were used for further tests.

**ChIP Assays**

Chromatin immunoprecipitation was performed according to previous reports (Liu et al., 2019; Zhu et al., 2012). In brief, 3 g of 0.5 cm^2^ fruit pieces of *SlBES1-OE-3* at mature green stage were collected and cross-linked using 1%(v/v) formaldehyde under vacuum for 10 min and ground to powder in liquid nitrogen. Then, the chromatin complexes were isolated as described previously (Liu et al., 2019), sonicated with Biorupter plus (Diagenode, Belgium, Antwerp), and immunoprecipitated with 5 μg Mouse monoclonal anti-c-myc antibody (clone 9E10, IgG, Roche, Basel, Switzerland). About 200~300 bp of ChIP DNA and input DNA was recovered and dissolved in water for further ChIP-qPCR analysis. For ChIP-qPCR, primer pairs were used to analyze the ChIP DNA (Supplemental Table 1). Each ChIP value was normalized to its respective input DNA value. The fold enrichment on each candidate genes or *PMEU1* promoter was calculated against the *ACTIN7* or *ACTIN2*, respectively. The mean value of two technical replicates was recorded for each biological replicate. The values of three independent biological replicates were collected and error bars represented the standard error from these tests.

**Transient Expression Assay in Tobacco Leaves**

Transient expression assay in tobacco (*Nicotiana benthamiana*) was performed as previously described (Shao et al., 2019). The *PMEU1* promoter was cloned with KOD FX (Toyobo, Japan, Osaka) and inserted into PQB using DNA Ligation Kit (Takara, Kusatsu, Japan). Then the promoter was fused with the LUC reporter gene into the plant binary vector pGWB35 using Gateway cloning kit (Thermo, Boston, USA) to generate the PMEU1*pro*:LUC reporter construct. *SlBES1*-1300 was used as an effector construct. These constructs were transformed into Agrobacterium cell GV3101 with P19 protein and pSoup vector, and then the cells were incubated, harvested and re-suspended in infiltration buffer (10 mM MES, 40 mM AS, and 10 mM MgCl_2_) to a final concentration of OD_600_= 1.0. Equal volumes of transformed cells with different combinations were mixed and then co-infiltrated into tobacco leaves with a needleless syringe. Infiltrated plants were placed at 28℃ for 48 h before imaging. NightOWL II LB983 Ultrasens backlit (Berthold technologies, Bad Wildbad, Germany) with Indigo software was used to capture LUC expression image and to quantify LUC luminescence intensity. 100 mM luciferin was sprayed on leaves and incubated in dark for 5 min before detection. Six independent determinations were performed.

**EMSA**

Full-length coding region of *SlBES1* was amplified and then inserted into *pET28a* vector. The recombinant protein, SlBES1-His were expressed in *E.coli* Rosetta cells at 28℃ and purified to homogeneity with Ni-His Resin (Thermo, Boston, USA) after identified by SDS-PAGE. Oligonucleotide probes were synthesized and labeled with FAM at 3’ ends. EMSAs (electrophoretic mobility shift assay) was performed as previously described (Liu et al., 2019) with minor modification. In brief, the FAM-labeled binding box probe, probe without any label (competitor) and mutant probe (mutant competitor) were incubated with SlBES1-His proteins at room temperature for 20 min, respectively. Bound and free probes were separated via PAGE. Typhon FLA7000 (GE, Fairfield, USA) was used to capture the final image. Probes were listed in Supplemental Table 2.

**Determination of Pectin Methylesterase Activity**

Cell wall extraction was conducted according to previous reports (Freitas et al., 2012). Alcohol-insoluble substances (AIS) from cell wall was first obtained and then soluble and insoluble pectin were extracted from AIS. 4 g of fresh tomato pericarp at different fruit stage was boiled in 25 mL 95% ethanol for 20 min and then mixed for 60s. The precipitate was left after centrifuged at 1500g for 10 min. Then the precipitate was mixed with 25 mL 80% ethanol and the mixture was centrifuged at 1500 g for 20 min to obtain precipitates. This step was repeated three times until the supernatant showed colorless. The crude cell wall pellet was dried under air steam and then suspended in DMSO: water (9:1, v/v, 20 mL/g). After stirring in room temperature for 24 h, the slurry was centrifuged at 1500 g for 20 min to remove DMSO. The pellet was washed repeatedly in 95% ethanol and then was dried under air. The dried material was washed once with acetone. Then this cell wall extraction was freeze-dried and weighted. It was centrifuged at 10500 g for 25 min after suspend in sterilized distilled water for 1h and this step was repeated for once. Supernatant and precipitate in these two steps were collected, respectively. The supernatant was soluble pectin while the precipitate was insoluble pectin. The pH was adjusted to 6 with 0.1M NaOH to ensure total ion amount of carboxyl group.

For pectin methylesterase (PME, EC3.1.1.11) activity determination, titration with an automatic titration was used as described in former reports (Tucker et al., 1992) with minor modification. Weighted crude protein from cell wall was assayed in determination solution, 2% (w/v) citrus pectin (pH=7.0). The volume of 5 mM NaOH consumed for titration was recorded over 5 min. Total PME activity was calculated as the volume of consumed NaOH per min (U·min^-1^·g^-1^).

**Degree of Pectin Methylesterification Determination**

The determination for methylesterifcation degree (DM) of pectin was performed as previously reported (Kyomugasho et al., 2015; Chylinska et al., 2016). Briefly, soluble and insoluble pectin was dissolved in deionized water. The pH was adjusted to 6 using 0.1M NaOH. After dialysis and freeze-drying, small pieces were compacted tightly to remove residual air and ensure smooth surfaces in the sampler of FT-IR (AVATAR 370 FT-IR, Thermo Nicolet, Boston, USA). 100 scans with 4000 cm^-1^ to 400 cm^-1^ transmittance wave numbers were run per sample to obtain spectra mean value with reduced noise. Therefore, DM could be calculated as the ratio of band area at 1749 cm^-1^ (represent as ester carbonyl group (C=O)) to the sum of band area at 1749 cm^-1^ and 1630 cm^-1^ (represent as carboxylate group (COO^-^)).

**Immunofluorescence**

Fresh tomato pericarp materials were fixed with FAA for at least 24h. Fixed tissue was dehydrated by serial incubations at 4℃ in solution with an increasing concentration of ethanol from 10% to 100%, and then submerged with LR White resin (Sigma-Aldrich, St. Louis, MO) to place into capsules.1 μm sections were obtained from the embedding block and then placed on glass slides for immunofluorescence assays. LM19 and LM20 mouse monoclonal antibodies bind demethylated and hypermethylated HGs, respectively. Additional, 2F4 mouse monoclonal antibodies are used to bind "egg box" epitopes formed of chain dimer of de-esterified HG linked with calcium ions (Silva-Sanzana et al., 2019). The non-specific binding sites were blocked by incubating the slide with the sample portion at room temperature for 30 min with 5% fat-free milk powder dissolved in 1 PBS and washed once with 1 PBS. The primary antibody was diluted with blocking solution (5%, 1×PBS) at a ratio of 1: 5, and the solution was incubated for 90 min at room temperature. Then samples were washed for three times with 1×PBS before secondary antibody incubation. The secondary antibody Alexa Fluor 488 goat anti-rabbit (Jackson ImmunoResearch, Pennsylvania, USA) was diluted with blocking solution (5%, 1×PBS) at a ratio of 1: 100 and incubated for 60 min at room temperature. Then 1× PBS solution was used to wash for three times. 0.25 mg / mL Calcofluor White (Sigma-Aldrich, St. Louis, MO) dissolved in 1×PBS was added for 5 min to fix the cell wall. This sample was washed with 1× PBS for twice and anti-fluorescence decay quencher Citifluor (Agar Scientific, Stansted, UK) was added before placing the coverslip. The images were observed with a NIKON ECLIPSE Ci-L upright microscope and an objective lens CFI 10*/22. The immunolabels of different treatments were done at least three times, and the most representative batch of processed images was selected for display. The relative signals of images were calculated through software ImageJ.
