## supplemental table3 for "Brassinosteroid signaling component SlBES1 promotes tomato fruit softening through transcriptional repression of *PMEU1*"

Supplemental Table S3. The contents of carotenoids and ascorbic acid in *SlBES1-OX* and *SlBES1-RNAi* fruits.

| Content  (μg/g FW) | lycopene | lutein | β-carotene | total carotenoid | ascorbic acid |
| --- | --- | --- | --- | --- | --- |
| AC | 164.41±11.95a | 6.52±0.10b | 117.46±4.81a | 245.93±38.20a | 180.91±14.03a |
| *SlBES1-OX-3* | 144.89±22.56a | 7.45±0.75a | 110.34±3.35a | 260.23±22.46a | 191.08±11.59a |
| *SlBES1-OX-8* | 148.04±20.99a | 7.42±0.34a | 108.13±5.50a | 263.55±27.86a | 193.98±5.60a |
| *SlBES1-RNAi-8* | 133.49±23.48a | 9.76±1.72a | 116.63±3.63a | 226.38±26.09a | 171.10±21.91a |
| *SlBES1-RNAi-9* | 127.51±29.62a | 8.02±0.47a | 114.49±3.62a | 250.02±30.57a | 168.20±15.31a |
| *SlBES1-KO* | 136.34±21.43a | 5.94±0.29b | 123.91±4.39a | 225.63±23.43a | 218.37±14.9a |

The contents of lycopene, lutein, β-carotene, total carotenoid and ascorbic acid in fruits of *SlBES1-OX* and *SlBES1-RNAi* at red ripe stage. Each data represents the means of three biological replicates with standard error. Different letters represent significant differences determined by Student’s t test (**p* < 0.05)
